# Tonic GABA_A_ receptor currents in Cerebellar Purkinje cells of wild-type and DMD^mdx^ mice

**DOI:** 10.1101/2025.07.07.663562

**Authors:** Shaarang Mitra, Shailesh N. Khatri, Jason R. Pugh

**Affiliations:** University of Texas Health Science Center at San Antonio, Department of Cellular and Integrative Physiology, San Antonio, TX 78229; Oklahoma State University Center for Health Sciences, National Center for Wellness and Recovery, Tulsa, Oklahoma, 74107-1898

## Abstract

Cerebellar Purkinje cells (PCs) fire spontaneously in the absence of excitatory input and depend heavily on inhibition to modify their firing activity. Previous work in the field has described phasic inhibition arising primarily from molecular layer interneuron–PC (MLI-PC) synapses extensively, however little work explores other sources of inhibition in PCs. Several types of neurons throughout the brain and within the cerebellum receive significant inhibition through tonic currents, a low amplitude current resulting from ambient GABA acting upon extrasynaptic GABA_A_ receptors. Through the use of ex vivo electrophysiology and single cell RNA analysis, we investigated the role of tonic inhibition in PCs. We find that PCs have a significant tonic current mediated by δ-subunit containing GABA_A_ receptors, which accounts for roughly half of the total inhibitory current. We also examined PC tonic GABA currents in DMD^mdx^ mice, a mouse model of Duchenne Muscular Dystrophy with ∼50% reduction in phasic inhibitory currents. We find that tonic inhibition is dramatically upregulated in DMD^mdx^ PCs, suggesting a possible compensatory mechanism to account for the loss in phasic inhibition. Furthermore, roughly 80% of the total inhibition is derived from tonic currents in this condition. These data suggest that under physiological conditions, PCs are subject to both tonic and phasic inhibition, and that adjustments in the balance of inhibition may be a physiological mechanism for PC function. These data reveal an expanded range of inhibitory currents in PC which may be critical to regulating PC activity in both normal and pathophysiological states.

## INTRODUCTION

Neuronal circuits require a precise balance of excitation and inhibition to properly encode and transmit signals. In the cerebellar circuit, Purkinje cells (PCs) receive excitation from parallel fibers and climbing fibers and inhibition primarily from molecular layer interneurons (MLIs). PC inhibition is primarily mediated by activation of GABA_A_ receptors, multimeric receptors generally consisting of 5 subunits: 2 α-subunits, 2 β-subunits, and either a γ- or δ-subunit. Functional GABA_A_ receptors can broadly be divided into synaptic receptors, those localized to the postsynaptic density, mediating phasic inhibition, and containing a γ-subunit; and extrasynaptic receptors, those localized outside the postsynaptic density, mediating tonic inhibition, and often containing a δ-subunit (Brickley and Mody, 2012; Luo et al., 2013). The synaptic γ-subunit containing GABA_A_ receptor (γ-GABA_A_ receptor) generally exhibit relatively low affinity for GABA (5-10 μM) and respond to high concentrations of GABA experienced at the synaptic cleft, whereas δ-subunit containing GABA_A_ receptor (δ-GABA_A_ receptor) generally have high affinity for GABA (<1 μM), enabling them to respond to relatively low levels of ambient GABA in the extracellular space (Mortensen et al., 2012; Zheleznova et al., 2009). In the cerebellar circuit, tonic inhibitory current are well described in granule cells, where they reduce noise, alter synaptic gain, maintain temporal fidelity of mossy fiber inputs, and participate in social and maternal behaviors (Chadderton et al., 2004; Duguid et al., 2012; Mitchell and Silver, 2003; Semyanov et al., 2004; Rudolph et al., 2020). Tonic GABA currents have also been described in PCs (Kueh et al., 2011), however, the molecular identity of the receptors involved or the physiological consequences of tonic currents have not been explored.

DMD^mdx^ mice are a model of Duchenne muscular dystrophy lacking expression of the protein dystrophin. In addition to muscle weakness and degeneration, humans with this disease and mouse models of the disease display a range of cognitive deficits and comorbidities with neurodevelopmental disorders (Ricotti et al., 2016; Fujino et al., 2018; Pane et al., 2012). Previous work has shown that dystrophin is also expressed in select regions of the brain, where it localizes to the postsynaptic density of inhibitory synapses, acting as a cell-adhesion molecule and/or participating in clustering GABA_A_ receptors (Anderson et al., 2003; Kueh et al., 2011). In cerebellar PCs, DMD^mdx^ mice show a significant reduction in both the number and strength of inhibitory synapses, dramatically reducing phasic inhibition (Wu et al., 2022). Interestingly, the rate of spontaneous action potential (AP) firing is unchanged, suggesting compensatory mechanisms to maintain overall inhibition and firing rate. One such mechanism may be upregulation of δ-GABA_A_ receptors and tonic GABA currents in PCs of DMD^mdx^ mice.

In this study we investigated whether PCs express δ-GABA_A_ receptors and exhibit a tonic inhibitory GABA current. We find that blocking GABA_A_ receptors reduces the holding current in PCs, indicative of an underlying tonic GABA current. This current persists in the presence of the voltage-gated sodium channel blocker, TTX, suggesting that ambient GABA near PCs is not dependent on AP generation or AP-evoked synaptic release of GABA. The tonic GABA current is enhanced by δ-subunit specific pharmacological agents, and abolished in a δ-subunit knockout mouse, suggesting the tonic current is mediated by δ-GABA_A_ receptors. Evoked firing in PCs is inhibited by pharmacological enhancement of the tonic GABA current, suggesting the tonic GABA current is able to modulate PC firing. In DMD^mdx^ mice, which show reduced phasic inhibition, we observed increased tonic GABA currents and increased sensitivity to low concentrations of GABA, consistent with increased expression of δ-GABA_A_ receptors. This is the first study showing the presence of δ-GABA_A_ receptor-mediated tonic current in cerebellar PCs. As cerebellar PCs are the sole output neurons of cerebellar cortex, we speculate that tonic GABA currents play a role in modulating PC excitability, noise reduction/normalization of incoming information from parallel fibers and PC related dysfunctions.

## METHODS

### Animals

All experimental procedures involving animals were approved by the Institutional Animal Care and Use Committee at UT Health San Antonio. Male and female C57BL/6 or Gabrd-KO (Mihalek et al., 1999; JAX stock #003725) juvenile (P14-P26) or adult (P50 and P90) mice (Charles River, MA) were used for tonic GABA experiments. P30-50 DMD^mdx^ mice (C57BL/10ScSn-Dmd*^mdx^*/J; catalog #001801, The Jackson Laboratory), a mouse model of Duchenne muscular dystrophy, were also used for tonic GABA current experiments. Only male DMD^mdx^ mice were used to better replicate the genetic background of Duchenne muscular dystrophy (an X-linked condition primarily affecting males). Animals were kept on a 12/12 hour light dark cycle with *ad libitum* access to food and water.

### Slice preparation

Mice were deeply anesthetized with isoflurane and the cerebellum was rapidly dissected following decapitation and placed in ice-cold oxygenated artificial cerebrospinal fluid (ACSF) containing the following (in mM): 119 NaCl, 26.2 NaHCO_3_, 2.5 KCl, 1.0 NaH_2_PO_4_, 11 glucose, 2 CaCl_2_, 1.3 MgCl_2_. Parasagittal slices (250-300 µm) were cut from the vermis of the cerebellum using a vibratome (Leica Biosystems, IL) and then incubated at 34°C for 30 min, after which they were kept at room temperature and used for electrophysiological recordings at room temperature.

### Electrophysiology experiments

For recording, slices were gently transferred, using a wide mouth pasture pipette, to a recording chamber that was perfused with ACSF containing the following (in mM): 119 NaCl, 26.2 NaHCO_3_, 2.5 KCl, 1.0 NaH_2_PO_4_, 11 glucose, 2 CaCl_2_, 1.3 MgCl_2_ (flow rate of ∼2 ml/min) housed in a SliceScope Pro upright microscope (Scientifica Instruments, UK). Voltage-clamp recordings were made from PCs using the following internal solution (in mM): 135 CsCl, 10 HEPES, 4 MgCl_2_, 5 EGTA, 4 Na-ATP, 0.5 Na-GTP, 2 QX-314. The pH of the internal solution was adjusted to 7.2-7.4 using CsOH and the osmolarity was 280-300 mosmol. Cells were patched using 2-4 MΩs borosilicate glass pipettes (Sutter instruments) that were pulled on a Sutter pipette puller (Model P-100, Sutter instruments, CA, USA). Electrophysiological currents were recorded with a Multiclamp 700B amplifier (Molecular Devices, CA), filtered at 5 kHz and digitized at 50 kHz. Data were collected using pCLAMP software (Molecular Devices). The tonic GABA_A_ receptor current was measured by the net change in holding current following application of 20 µM bicuculline (Tocris, MN). For all tonic GABA current recordings 10 µM NBQX and 10 µM CPP were added to the ACSF to block fast excitatory synaptic transmission. Where indicated, the ACSF also contained one or more of the following: 20 µM Bicuculline (Tocris), 20 µM DS2 (Tocris), 25 µM Gabazine (SR-95531, Tocris), 500 nm THIP (Tocris). Access resistances of all cells were monitored and cells deviating from 20±10 MΩs were not included.

### Tonic current and charge transfer analysis

To identify tonic current magnitude, we measured the average holding current 30 seconds immediately before and after drug wash. The average holding current was determined by creating a histogram of all data points during the 30 second period and fitting the histogram with a Gaussian function. The peak of the Gaussian fit was used as the average holding current and the tonic current was defined as the difference between the peak of the Gaussian pre- and post-drug application. Recordings without a stable holding current prior to drug application were discarded. To calculate total and phasic inhibitory charge transfer, we calculated the area under the curve to either the average baseline holding current (phasic charge transfer) or to the average holding current following drug application (total charge transfer). Charge transfer due to tonic GABA current was calculated by subtracting the phasic charge transfer from the total.

For current-clamp recordings, the internal solution contained (in mM) 137 K-gluconate, 4 KCl, 10 HEPES, 4 MgCl2, 5 EGTA, 4 Na-ATP, 0.5 Na-GTP (pH 7.3-7.4, 285-295 mosm). In these recordings, excitatory synaptic transmission was blocked by 10 μM CPP (Abcam) and 10 μM NBQX (Sigma) in the bath ACSF. Action potentials were evoked by depolarizing current injections (200 ms, 450 pA) through the patch pipette from a hyperpolarized baseline membrane potential (-70 mV). For each cell values were averaged across 3 sweeps.

### GABA Uncaging

For uncaging experiments, the internal solution was identical to the solution detailed above for voltage clamp recordings. In addition to the 10 µM NBQX and 10 µM CPP, 60 µM Rubi-GABA (Tocris) was added to the ACSF. A 473 nm wavelength blue light laser (Opto Engine LLC) was routed through the microscope objective and centered on the soma of the patched PC. Cells were exposed to increasing intensities of light, liberating increasing concentrations of GABA, generating a current in the patched cell. Each trial consisted of 2 laser pulses with an interstimulus interval of 1 s, repeated 5 times at each laser power. A 30 second waiting period was used between trials to allow for cell recovery and diffusion of previously liberated GABA. For each cell, the order of laser powers tested was randomized.

### Single cell patch qRT-PCR

Cytoplasmic RNA was collected through a whole-cell patch clamp pipette by applying light suction until the cell cytoplasm was aspirated (Citri et al., 2011; Cadwell et al., 2015; Fuzik et al., 2016). Patch pipettes were then gradually raised above the slice, occasionally removing the entire cell from the slice. Patch pipettes with nonspecific cell debris stuck from surrounding cells were discarded. The contents of the pipette were collected using positive pressure into an RNase free PCR tube containing lysis buffer. These cells were immediately transferred and stored at −80°C until further processing for single-cell qPCR. Single-cell gene expression analysis was performed using the Bioanalytic and Single-cell core (BASiC) at UT Health at San Antonio as described previously (Chen et al., 2013). The BASiC core is supported by the Cancer Prevention Research Institute of Texas (RP150600) and the Office of Vice President of Research of UT Health at San Antonio.

Briefly, single-cell patch qRT-PCR was carried out using Cells Direct^TM^ one-step RT-PCR kit (11753–100, Invitrogen, Waltham, MA, USA) and a microfluidics device, BioMark HD MX/HX system (cat #BMKHDPKG-MH, Fluidigm, Inc., South San Francisco, CA). The cytoplasm from single cells in lysis buffer were thawed, mixed well, and spun down before being lysed at 75°C for 10 min. Genomic DNA contamination was reduced by treating cells with DNase I. PCR primers of selected genes were divided into two panels to fit BioMark 12 × 12 or 48 × 48 chips. Reverse transcription (RT), preamplification, and PCR amplification were carried out according to the protocol of single-cell gene expression (cat #BMK-M-12.12, Fluidigm). Target genes were amplified using BioMark HB MX/HX system with 1X SsoFast Eva-Green supermix with low ROX (172-5211, Bio-Rad, Hercules, CA) and 1X DNA binding dye sample loading reagent (100-3738, Fluidigm). In each chip assay, universal mouse RNA (200 pg) from mouse normal tissues (R4334566-1, BioChain, Newark, CA) and no template control (NTC) served as positive and negative controls. Quantitative PCR products were detected using Fluidigm BioMark HD system according to the protocol: Gene expression with the FlexSix IFC using Delta Gene assays (100-7717 B1, Fluidigm). GE Flex Six Fast PCR+Melt v1 program was used to collect the cycle threshold (CT) values. Raw CT values were obtained from the Fluidigm Biomark software and inverted (25-CT) to generate a log2-based scale for gene expression analysis and presentation.

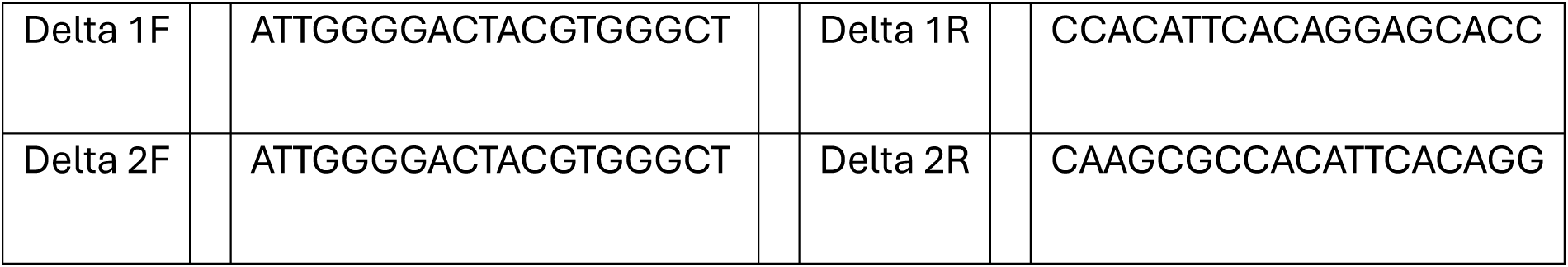

#### Data analysis

Data was analyzed in IgorPro (Wavemetrics, Lake Oswego, OR) using the Neuromatic toolkit (Rothman and Silver 2018) and custom macros. Statistical significance was determined using two-tailed paired or unpaired Student’s *t*-tests in Excel (Microsoft, Redmond, WA) or Prism 6 (GraphPad Software, La Jolla, CA, USA). Statistical values of *P* ≤ 0.05 were considered significant.

## RESULTS

### Cerebellar PCs exhibit GABA_A_R mediated tonic currents

To examine tonic GABA currents in cerebellar PCs, we used whole-cell patch clamp electrophysiology in acute cerebellar slices from juvenile C57B/L6 mice. Upon application of GABA_A_ receptor antagonists (20 µM bicuculline or 25 µM gabazine) spontaneous IPSCs were rapidly blocked. In addition, we observed a reduction in the holding current (figure 1A), indicative of blocking a tonic GABA_A_ receptor mediated current. On average, the tonic GABA current was 24.8±3.9 pA in bicuculline (n=19) and 20.3±5.3 pA in gabazine (n=6, figure 1C). To determine whether this current persists beyond developmental stages, we repeated this experiment in adult mice (P50 and P90). At both adult ages, we observed a reduction in PC holding current following application of bicuculline (P50: 17.2±34.9 pA, n=6; P90: 9.1±2.9 pA, n=6, figure 1C), suggesting adult PCs also express a tonic GABA current, however the magnitude of the current decreased with age (P15 vs P90, p=0.04). In the granule cell layer, it has been proposed that ambient GABA results from AP-dependent synaptic GABA release (Brickley et al., 1996; Wall and Usowicz, 1997) or through AP-independent GABA release from astrocytes and glia (Lee et al., 2010). To test these possibilities in PCs, we preincubated slices in 500 nM TTX, to block AP generation, prior to application of bicuculline. The presence of TTX did not reduce the amplitude of tonic GABA currents in PCs (19.9±3.1 pA, n=8, p=0.46, figure 1B, C), suggesting that the GABA responsible for the tonic currents is not mediated by AP-dependent GABA release. Rather, ambient GABA likely arises from either spontaneous vesicle fusion at inhibitory terminals (miniature IPSC were prominent in our PC recordings in TTX; figure 1B) or by release from Bergmann glia (Barakat and Bordey 2002). Together, these data show that PCs exhibit an AP-independent tonic GABA current at both juvenile and adult ages.

**Figure 1.**
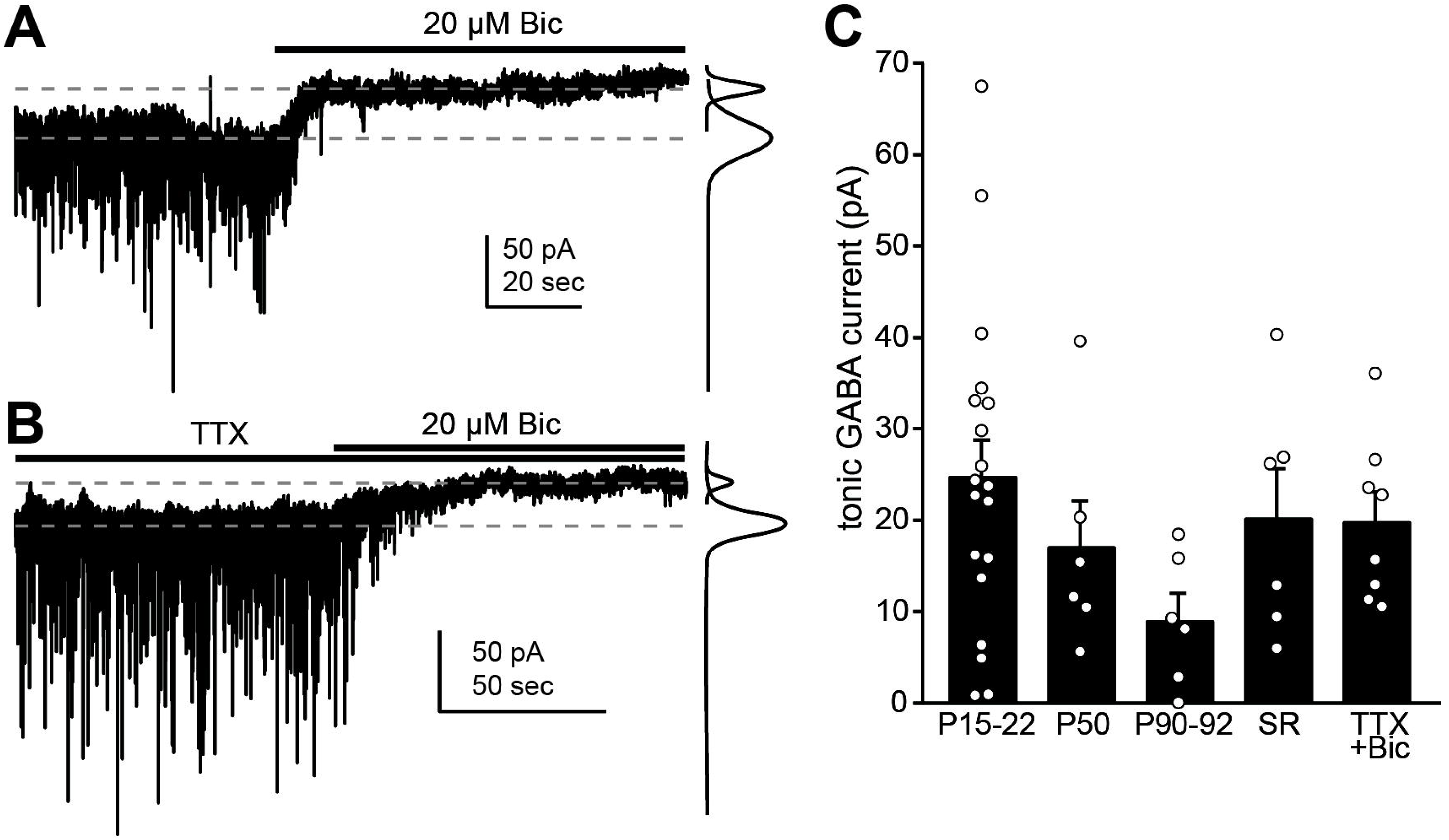
Measurement of tonic GABA currents in PCs. A) Representative current-trace of PC held at -60 mV. Tonic GABA currents are calculated by the difference in average holding current before and after bicuculline application (dashed lines). Gaussian fits of histograms of current values in each condition are shown on the right. B) Same as in (A) but recorded in the presence of 500 nM TTX. C) Average tonic GABA_A_ currents amplitudes across age ranges (P15-22, P50, and P90-92). Tonic current amplitude decreases with age (p=0.04). Average tonic GABA_A_ currents amplitudes following application of gabazine or in the presence of TTX is not different from currents measured with bicuculline at the same age. Values from individual cells are plotted as open circles.

### Tonic currents in cerebellar PCs are dependent on δ-GABA_A_ receptors

In cerebellar granule cells, the tonic GABA current is mediated by extrasynaptic δ-subunit containing GABA_A_ receptors (Hamann et al., 2002). In order to determine whether this is also true of tonic GABA currents in PCs, we first examined if PCs encode for δ-subunits. We extracted cytoplasmic material from WT PCs, along with cerebellar granule cells and HEK cells or oligodendrocytes to act as positive and negative controls. Utilizing qRT-PCR, we amplified δ-subunit coding mRNA, and observed robust expression in PCs and GCs, but not HEK cells or oligodendrocytes (figure 2A). mRNA levels were not significantly different between GCs and PCs (p=0.48), suggesting PCs code for significant levels of δ-subunits. Furthermore, δ-subunit mRNA expression was significantly reduced in δKO PCs (p<0.05). Together, these data suggest PCs express δ-subunit mRNA.

**Figure 2.**
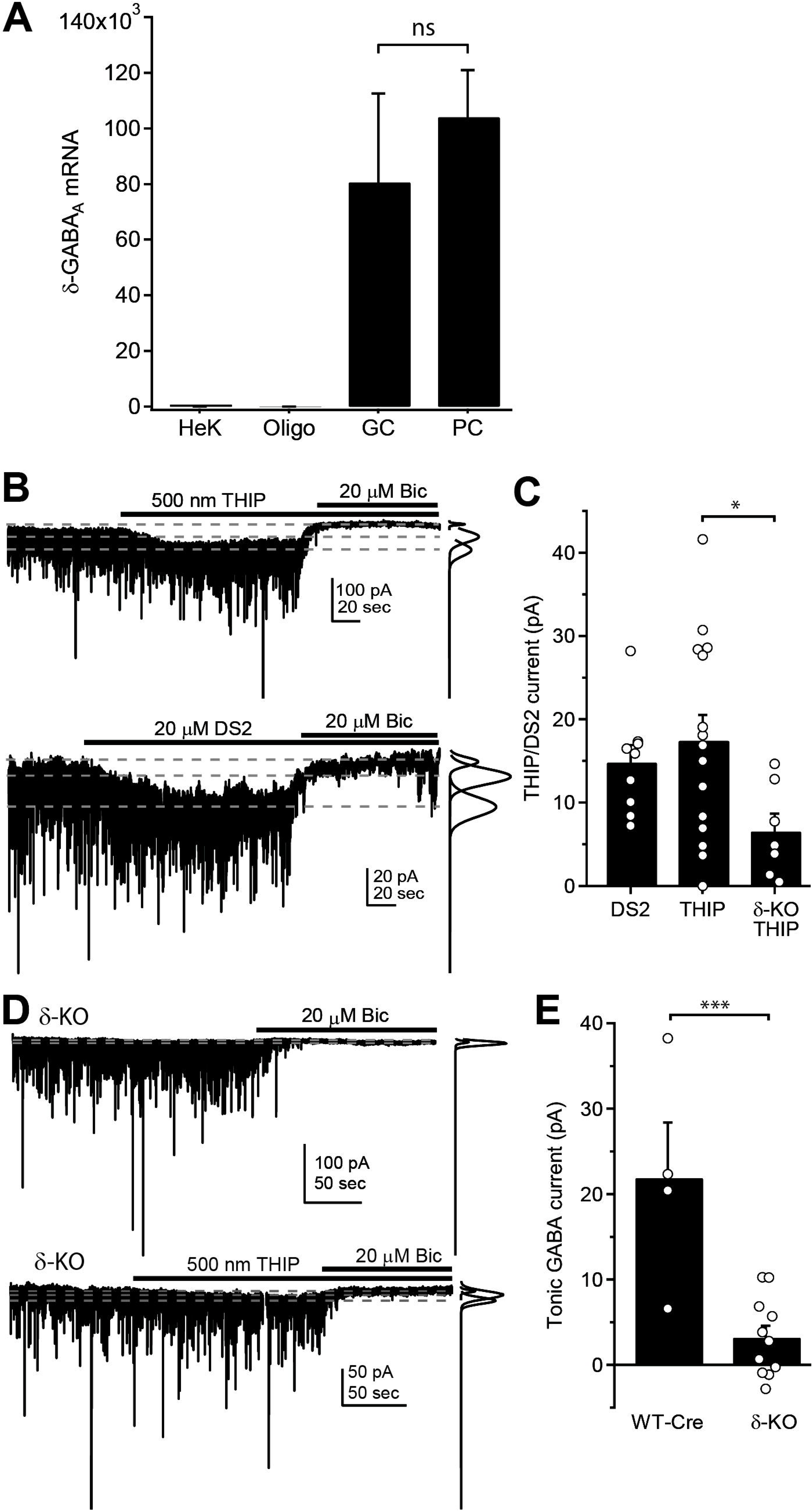
δ-subunit containing GABA_A_ receptors contribute to tonic GABA currents in PCs. A) GABA_A_ δ-subunit mRNA expression measured by single cell qRT-PCR in HeK cells, oligodendrocytes, cerebellar granule cells, and Purkinje cells. B) THIP, a δ-subunit specific agonist (top), and DS2, a δ-subunit specific positive allosteric modulator (bottom), shift the holding current in PCs by enhancing tonic GABA currents. C) Average increase in PC holding current following application of THIP or DS2. D) Current traces showing lack of tonic GABA current (top) and THIP current (bottom) in δ-KO PCs. E) Average tonic GABA current recorded in PCs from δ-KO or WT littermate controls. Values from individual cells are plotted as open circles. Gaussian fits of histograms of current values in each condition are shown on the right (B and D).

To assess expression of functional δ-GABA_A_ receptors, we measured changes in the PC holding current following application of either, THIP, a δ-GABA_A_ receptor specific agonist at low concentrations (Krogsgaard-Larsen 1984, Drasbek 2006) or DS2, a δ-GABA_A_ receptor positive allosteric modulator (PAM; Wafford 2009). We observed an increase in the holding current of PCs upon bath application of either THIP (17.4±3.1 pA, n=15) or DS2 (14.8±2.1 pA, n=9), consistent with expression of δ-GABA_A_ receptors (figure 2B, C). In order to confirm that currents induced by THIP or DS2 are mediated by GABA_A_ receptors, we applied bicuculline (20 µM) at the end of each experiment. Following application of bicuculline, the holding current was consistently reduced below the baseline prior to THIP/DS2 administration (figure 2B). The tonic GABA current measured in bicuculline (relative to the baseline before THIP/DS2 administration) was not different from tonic GABA currents measured in the absence of either drug (p>0.05) suggesting that currents induced by either THIP or DS2 are mediated by GABA_A_ receptors. These data suggest that GABA_A_ receptors containing a δ-subunit contribute to tonic GABA currents in cerebellar PCs.

In order to further demonstrate the δ-subunits containing receptors contribute to tonic GABA currents in PCs and determine whether other types of GABA_A_ receptors may also contribute, we measured tonic GABA currents in PCs from a δ-subunit knockout mouse line (δ-KO). In δ-KO PCs we observed little or no change in holding current following application of bicuculline (3.2±1.4 pA, n=11, figure 2D, E), a significant reduction from tonic GABA currents observed in PCs of WT littermate controls (21.9±6.5 pA, n=4, p=0.0008) or c57B6 animals (p=0.0004, figure 1C). Furthermore, bath application of THIP produced little change in the holding current of δ-KO PCs (6.5±2.1 pA, n=7, figure 2C,D). These data suggest that tonic GABA current in PCs are primarily mediated by δ-GABA_A_ receptors.

### δ-GABA_A_ receptors modulate excitability of cerebellar PCs

PCs are relatively large, spontaneously active neurons with substantial Na^+^ and K^+^ currents during AP firing. However, tonic GABA currents in these cells are relatively small (∼20 pA), raising the question of how much these currents affect PC activity. To test this, we made current injections (450 pA) into PCs to evoke firing from a hyperpolarized baseline before and after application of THIP. We found that the number of action potential evoked by the current injection was reduced by application of THIP (cnt: 11.6±2 APs; THIP: 9.82±1.6 APs; n=9, p=0.03 paired t-test; figure 3A,B), suggesting modulation of tonic GABA currents is sufficient to reduce PC activity.

**Figure 3.**
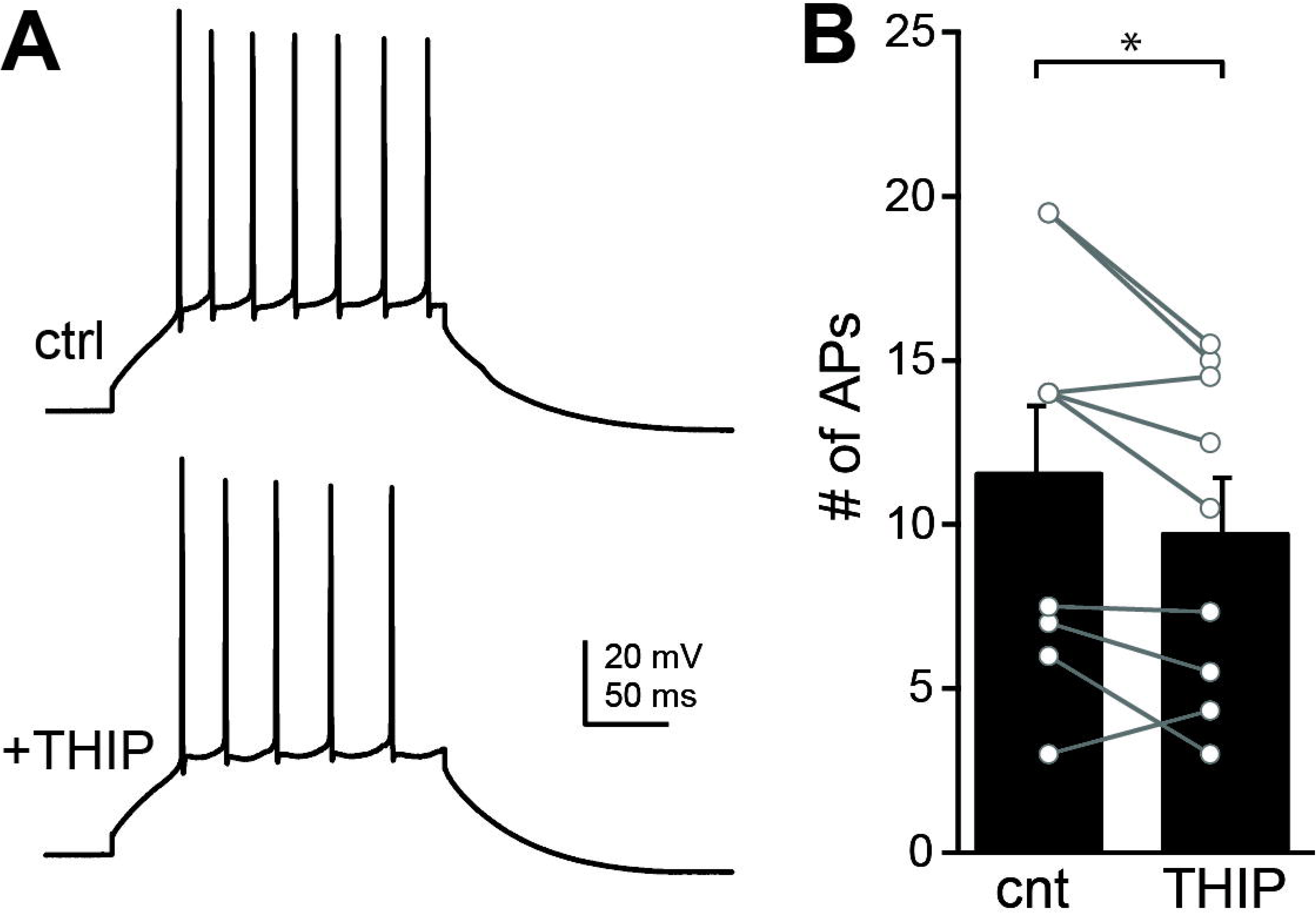
δ-subunit-mediated currents influence PC firing. A) Example voltage traces of PC firing in response to current injection in the same cell before (top) and after (bottom) application of THIP. B) The average number of action potentials during the current injection is reduced by THIP. Values from the same cell are plotted as connected open circles.

### DMD^mdx^ mice express larger tonic GABA currents in PCs

DMD^mdx^ mice, a model of Duchenne muscular dystrophy (DMD) lacking expression of the protein dystrophin, display an approximately 50% reduction in inhibitory synapses number and phasic inhibition in cerebellar PCs (Wu et al., 2022). This raises the possibility that tonic GABA currents are increased in DMD^mdx^ PCs, either in response to reduced phasic inhibition or as a result of reduced GABA_A_ receptor clustering at postsynaptic densities (Kueh et al., 2011). To test this possibility, we recorded tonic GABA currents from male DMD^mdx^ mice and WT littermate controls. Though tonic GABA currents were generally smaller in the DMD^mdx^ background, we found a three-fold increase in tonic current in DMD^mdx^ PCs (21.5±2.0 pA, n=6) compared to WT littermate controls (6.1±1.8 pA, n=10 cells, p<0.0001, figure 4A-C). While the magnitude of these currents is relatively small, their tonic nature can result in large changes in charge transfer across the membrane and total inhibition. We found that total charge transfer over a 30 second interval (due to both phasic and tonic GABA inhibition) was significantly increased in DMD^mdx^ PCs (758.8±82.0 pC, n=6) compared to WT (273.1±52.3 pC, n=10, p=0.0001, figure 4D) despite the reduction in phasic currents in DMD^mdx^ PCs. This increase was due to a substantial increase in inhibitory charge transfer due to tonic GABA currents in DMD^mdx^ PCs (192.2±61.6 pC vs 707.1±125.8 pC, p=0.001). Further analysis showed that the percent of total inhibitory charge transfer mediated by tonic currents is increased in DMD^mdx^ PCs compared to WT PCs (WT:53.9±11.0%, DMD^mdx^:89.2±2.4%, p=0.03, figure 4D). These results show that tonic GABA currents can account for a large fraction of total inhibition, and that increased tonic currents in DMD^mdx^ PCs may compensate for loss of phasic inhibition.

**Figure 4.**
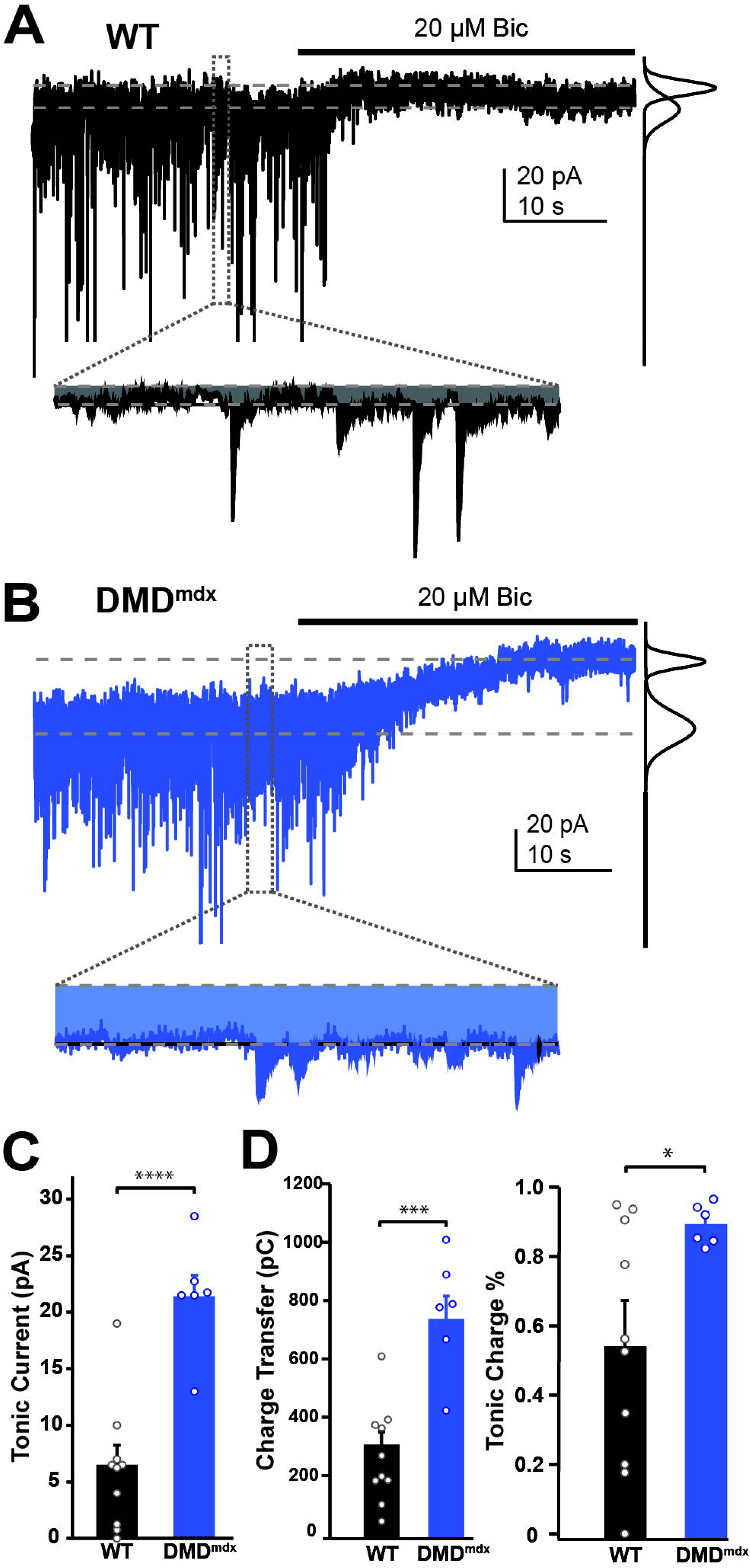
Tonic GABA currents are increased in DMD^mdx^ mice compared to WT littermate controls. Representative current traces of tonic GABA currents in PCs from DMD^mdx^ mice (B, blue) and WT littermate controls (A, black). Dashed lines represent average holding current before and after bicuculline application. Gaussian fits of histograms of current values in each condition are shown on the right. Lower traces are expanded regions of each trace showing inhibitory charge transfer (area under the curve) due to phasic inhibition (black/dark blue shaded areas) or tonic inhibition (grey/light blue shaded areas). C) Average tonic GABA current in WT and DMD^mdx^ PCs. D) Total inhibitory charge transfer (left) and percent of total charge transfer due to tonic GABA currents (right) in WT and DMD^mdx^ PCs. Values from individual cells are plotted as open circles.

### δ-GABA_A_ receptor expression is increased DMD^mdx^ PCs

An increase in tonic GABA currents in DMD^mdx^ PCs could be due to reduced clustering of synaptic GABA_A_ receptors at synaptic densities, resulting in an increased number of extrasynaptic receptors able to respond to ambient GABA, or due to an upregulation of higher affinity δ-GABA_A_ receptors in response to reduced phasic inhibition. To test for increased δ-GABA_A_ receptor expression, we first recorded tonic currents from DMD^mdx^ and WT littermate PCs in the presence of DS2. DS2 had a more pronounced effect on tonic GABA currents in DMD^mdx^ PCs (p=0.02, figure 5B), compared to WT littermate controls (p=0.2). Additionally, DS2 tonic currents were significantly larger in DMD^mdx^ PCs compared to littermate controls (p=0.002). These data suggest that δ-GABA_A_ receptor expression is increased in DMD^mdx^ PCs, potentially accounting for the increased tonic currents.

**Figure 5.**
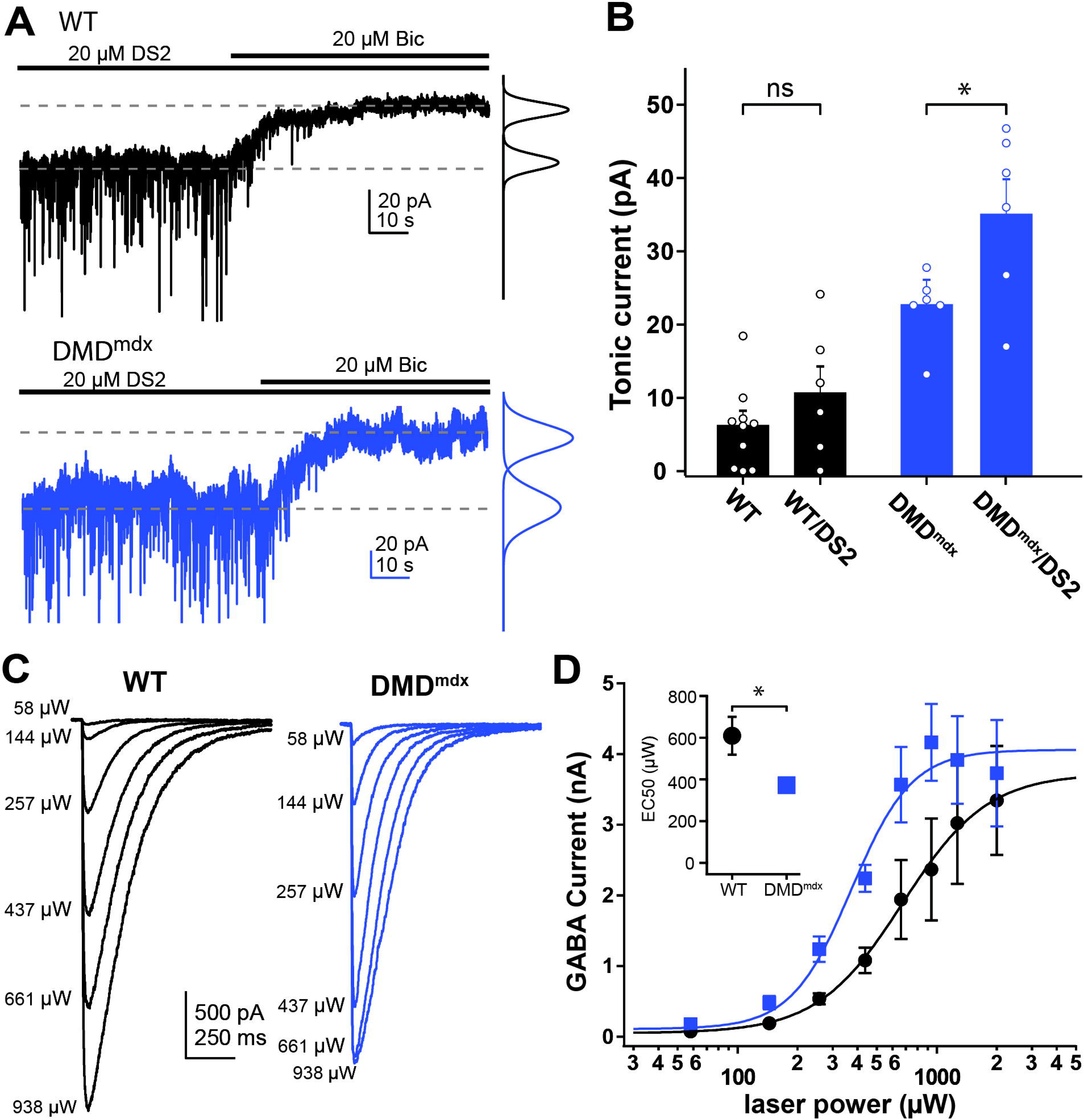
Tonic GABA currents in DMD^mdx^ PCs are mediated by expression of δ-subunit containing GABA_A_ receptors. A) Representative current traces showing tonic GABA current measured in the presence of DS2 from WT or DMD^mdx^ PC. B) Average tonic GABA current measured in control solution or in the presence of DS2 in WT (black) and DMD^mdx^ (blue) PCs. Values from individual cells are plotted as open circles. C) Representative traces of currents evoked by photolytic uncaging of Rubi-GABA over the soma of WT and DMD^mdx^ PCs using a range of laser powers (58-938 µW). D) Average GABA current response plotted against uncaging laser power for WT (black) and DMD^mdx^ (blue) PCs. Inset: Average EC_50_ value for WT and DMD^mdx^ derived from fitting the power-response curve for each cell with the Hill equation.

To further confirm upregulation of δ-GABA_A_ receptors, we measured the dose-response relationship of GABA_A_ receptors expressed on PCs in each genotype. Varying concentrations of GABA were applied by photolytic uncaging of RuBi-GABA by a 473 nm laser. We were able to liberate escalating concentrations of GABA at the PC soma by modulating the power of the laser, as we have done previously (Khatri et al., 2019). GABA uncaging was confined to a relatively small area (∼30 um) around the soma (Supplemental figure 1) and GABA currents in PCs varied with uncaging laser power (figure 5C). We found that GABA-evoked currents were larger in DMD^mdx^ PCs at low laser powers (58-437 μW p=0.002-0.03) but not at high laser powers (938-2000 μW p=0.1-0.77), producing a leftward shift in the power-response curve (figure 5D). By fitting the power response curves with the Hill equation, we find a significant reduction of the EC_50_ in DMD^mdx^ PCs (WT: 646.4±105.7 µW, n=5; DMD^mdx^: 372.1±33.7 µW, n=7; p = 0.017; figure 5D inset), but no change in the Hill coefficient (p=0.28) or maximum response (p=0.38). These data suggest an increase in the sensitivity of GABA_A_ receptors, with little change in the total number, consistent with increased expression of high-affinity δ-GABA_A_ receptors.

## Discussions

Neuronal activity is regulated by a dynamic balance of excitation and inhibition. While almost all neurons experience rapid phasic inhibition, driven by synaptically released GABA and activation of GABA_A_ receptors in the postsynaptic density, a handful of neurons also experience tonic inhibition, driven by ambient GABA activating extrasynaptic GABA_A_ receptors. In these neurons, tonic inhibitory currents can account for a large fraction of the total inhibitory charge transfer and significantly alter neuronal firing (Lee and Maguire, 2014; O’Neill and Sylantyev, 2018). We show that PC of the cerebellum also display tonic inhibitory currents revealed by a shift in the holding current following application of GABA_A_ receptor antagonists. Several lines of evidence suggest tonic GABA currents in PCs are primarily mediated by δ-GABA_A_ receptors, including qRT-PCR of δ-subunit mRNA from PC cytoplasmic extracts, enhancement of tonic currents by δ-subunit specific pharmacology, and loss of tonic currents in δ-subunit KO mice. Though tonic GABA currents are relatively small in PCs, pharmacological enhancement of these currents was sufficient to reduce evoked firing. To test the importance of tonic currents further, we measured these currents in DMD^mdx^ mice, which show a ∼50% reduction in inhibitory synapse number and phasic inhibition (Wu et al., 2022). We found the tonic GABA current is significantly increased in DMD^mdx^ PCs, potentially compensating for the loss of phasic inhibition and normalizing PC firing to the optimal range. In fact, the total inhibitory charge transfer is greater in DMD^mdx^ PCs compared to WT, despite the decrease in phasic inhibitory currents. Using photolytic uncaging of RuBi-GABA across a range of laser powers, we found a leftward shift in the dose-response curve with no change in the maximal current, suggesting the enhanced tonic current is associated with high-affinity δ-GABA_A_ receptor expression. Together, these data demonstrate a previously undescribed form of inhibition in cerebellar PC capable of modulating firing and compensating for loss of phasic inhibition in a disease model.

Phasic inhibition is generally mediated by synaptic GABA release activating GABA_A_ receptors clustered in the adjacent postsynaptic density. However, several neurons, including those in the hippocampus, thalamus, hypothalamus, brain stem, prefrontal cortex, cerebellum, and olfactory bulb also display a tonic GABA current mediated by extrasynaptic receptors (Farrant and Nusser, 2005; Brickley and Mody, 2012). Previous studies in the cerebellar circuit have focused on tonic GABA currents in granule cells mediated by α6δ-subunit containing receptors. In these cells, tonic GABA currents maintain a low firing rate (Chadderton et al., 2004), enhance the fidelity of sensory information (Duguid et al., 2012), modulate gain control during synaptic excitation (Mitchell and Silver, 2003), and participate in social and maternal behaviors (Rudolph et al., 2020). Our data show that tonic GABA currents in PCs can modulate the frequency of evoked APs and compensate for loss of phasic inhibition. Previous studies have shown that inhibition regulates calcium influx during climbing fiber activity and induction of synaptic plasticity at parallel fiber-PC synapses, a critical site for cerebellar learning (Gaffield et al., 2019; Zhang et al., 2023). Future studies, ideally employing PC specific δ-subunit KOs, will be required to determine the extent to which tonic GABA currents regulate these processes and higher circuit functions in the cerebellum.

Tonic GABA currents in cerebellar granule cells, thalamic neurons, and hippocampal neurons are associated with δ-GABA_A_ receptors (Zheleznova et al., 2009), however, other subunit combinations have also been shown to mediate tonic currents, for example, α5-containing receptors in CA1 pyramidal neurons (Cariascos 2004, Donegan 2019). Previous studies using IHC to label δ-subunits in the cerebellar cortex have observed strong labelling in the granule cell layer with only faint labelling in PCs (Pirker et al., 2000; Rudolph et al., 2020), leading to the conclusion that δ-subunits are exclusively expression in granule cells. Our data, on the other hand, suggests that tonic currents in PCs are primarily mediated by δ-GABA_A_ receptors. First, we observed δ-subunit mRNA in PCs using single cell cytoplasmic extracts, a method that can be more sensitive to low levels of expression compared to IHC. Second, tonic GABA currents were enhanced by application of either THIP, a δ-subunit specific agonist at low concentrations, or DS2, a δ-subunit specific PAM. Third, tonic currents were significantly reduced or absent in PCs from δ-subunit KO mice.

We speculate that weak IHC labelling in the PC or molecular layers may have been overlooked compared to the high cellular density and strong labelling in the granule cell layer. This difficulty is magnified by the difference in GABA_A_ receptor surface expression density between granule cells and PCs. Tonic currents in PCs are ∼20 pA, similar to currents observed in granule cells (Hamann et al., 2002; Rossi et al., 2003). However, the soma and dendritic arbor of PCs are much larger than granule cells (the PC soma alone is ∼5 times the diameter of a granule cells, translating to ∼25-fold increase in surface area), suggesting a similar number of GABA_A_ receptors are spread over a much greater (25-40x) surface area in PCs. This difference in channel density likely accounts for the lack of δ-subunit labelling in PCs using IHC.

In order to understand the physiological relevance of tonic currents in PCs, we also examined DMD^mdx^ mice, a model of Duchenne muscular dystrophy lacking dystrophin expression. In PCs, dystrophin is localized to inhibitory synapses where it acts as a transynaptic cell adhesion molecule (Briatore et al., 2020) and is involved in GABA_A_ receptor clustering (Knuesel et al. 1999, Grady et al., 2006). Our previous work has shown a ∼50% decrease in the number of inhibitory synapses and phasic inhibition in DMD^mdx^ PCs. Though tonic current amplitudes were generally lower in the DMD^mdx^ background, we found that currents were ∼3-fold higher in DMD^mdx^ compared to WT littermate controls. In fact, when measuring the total inhibitory charge transfer, we found greater inhibition in DMD^mdx^ compared to WT, suggesting that even small changes in tonic current amplitude (∼20 pA) can produce large changes in overall inhibition. Earlier studies have found that the total number of GABA_A_ receptors is not increased in the DMD^mdx^ brain (Zarrouki et al., 2022), including in the cerebellum specifically, suggesting the increased tonic current may be due to changes in either receptor localization or subunit composition. Previous studies have observed reduced GABA_A_ receptor clustering at postsynaptic densities in DMD^mdx^ PC (Knuesel et al., 1999; Kueh et al., 2011), suggesting synaptic GABA_A_ receptors may be mislocalized to the extrasynaptic space. Alternatively, a change in expression to higher affinity δ-GABA_A_ receptors could also increase the tonic current without a change in overall receptor number. Our data are more consistent with a change in subunit expression. We found that tonic currents in DMD^mdx^ are more sensitive to a δ-specific PAM, DS2, consistent with an earlier study showing enhanced sensitivity to THIP (Kueh et al., 2011). Furthermore, we observed a leftward shift in the dose-response curve and decrease in the EC_50_ of PC GABA_A_ receptors, consistent with increased expression of high affinity δ-GABA_A_ receptors. Earlier work measuring GABA_A_ receptor subunit expression by western blot analysis found a roughly 30% increase in δ-subunit expression in cerebellum of DMD^mdx^ mice, however this change did not reach significance (Zarrouki et al., 2022). Assuming little change in δ-subunit expression in granule cells, which do not normally express dystrophin, it is possible that even relatively large changes in δ-subunit expression in PCs would produce only a small change in total protein in cerebellar tissue measured by western blot analysis given the high density and number of granule cells compared to PCs, potentially accounting for the lack of significant increase in δ-subunit expression by western blot.

Tonic and phasic GABA currents likely play unique roles in sculpting PC behavior. Tonic currents are well-suited to regulate the average spontaneous firing rate of PCs, keeping firing in an optimal range. On the other hand, phasic currents arising primarily from molecular layer interneurons can pause PC firing and synchronize activity across PCs (Hausser and Clark, 1997; Lackey et al., 2024). Interestingly, we have previously shown that the average firing rate of DMD^mdx^ PCs is unchanged either in vivo or in ex vivo slices (Wu et al., 2022), likely due to the increase in tonic GABA currents compensating for loss of phasic inhibition. However, we also found that DMD^mdx^ PCs display significantly more regular spontaneous firing in ex vivo slices compared to WT, suggesting that while increased tonic current may restore total inhibition and the average firing rate, it does not mimic the stochastic nature of phasic inhibition. The resulting increase in regularity of firing may increase synchrony of firing across PCs, reduce the magnitude or frequency of bursts or pauses in PC firing, and alter signaling in downstream neurons (Luque et al., 2019; Person and Raman, 2011). This suggests that tonic and phasic currents play unique roles in determining PC activity. Future experiments using subunit specific pharmacology and/or genetic manipulations will be necessary to determine the extent to which phasic and tonic GABA currents regulate PC activity and plasticity.

## Acknowledgments

We are grateful to members of the Pugh lab for helpful discussion and comments. Single cell mRNA expression analysis was performed with the help of the Bioanalytic and Single-cell core (BASiC) at UT Health at San Antonio. This work was funded by National Institutes of Health, Grant # R01 NS123933.

## Figure Captions

**Supplemental figure – Spatial specificity of photolytic GABA uncaging.** A) Diagram showing locations of 488 nm laser pulses relative to the Purkinje soma. B) Average normalized current amplitudes and example traces (inset) following photolytic uncaging of RuBi-GABA over the soma (0 μm) and increasing distances (15-45 μm) from the center of the soma. Uncaging evoked-currents diminished with a space constant of 18.4 μm as the center of the uncaging laser was moved away from the soma.

