## Supplementary figures and images for "Tonic GABA_A_ receptor currents in Cerebellar Purkinje cells of wild-type and DMD^mdx^ mice"

### Supplemental figure 1

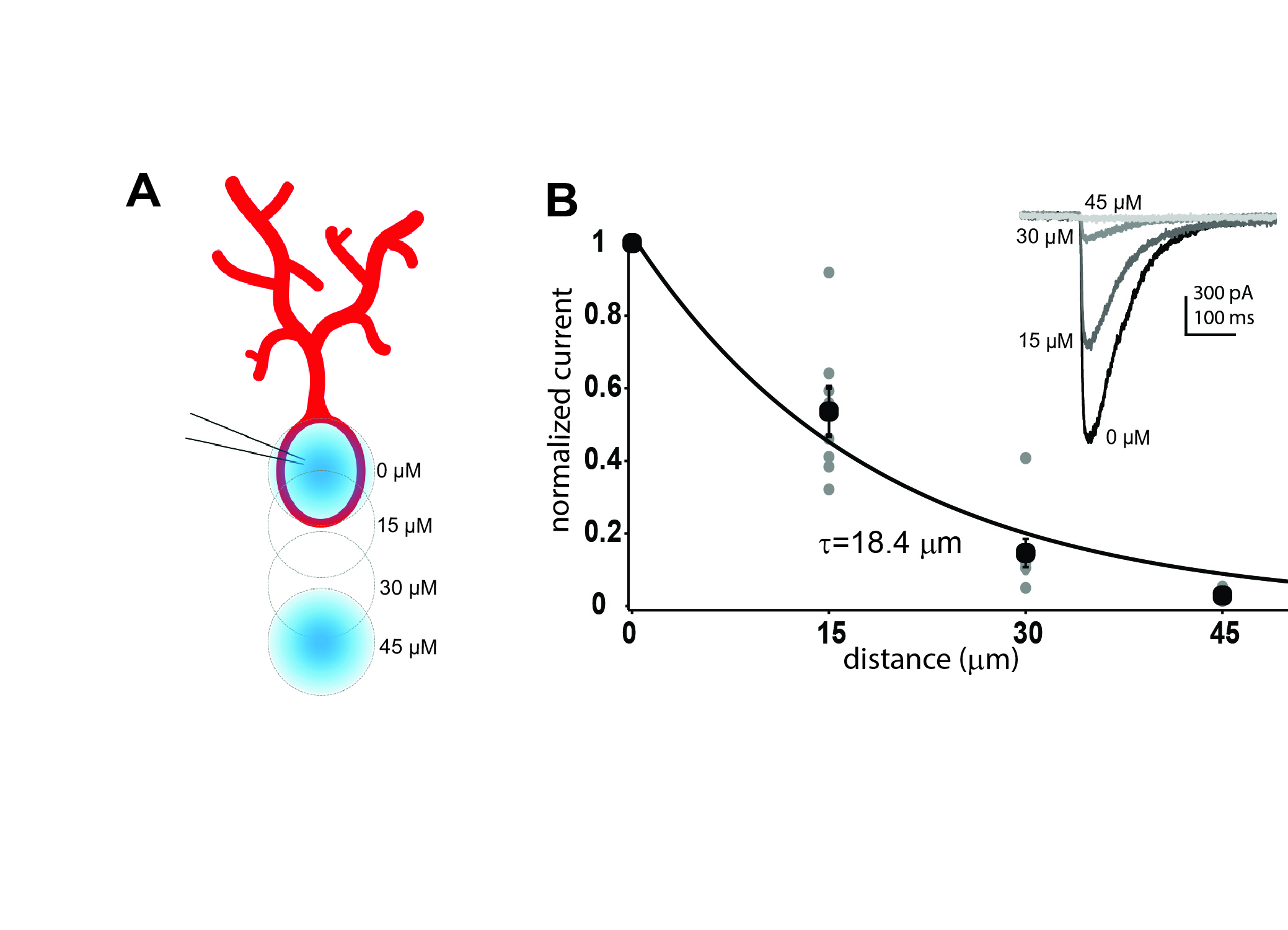
